# Identifying common transcriptome signatures of cancer by interpreting deep learning models

**DOI:** 10.1101/2021.11.11.467790

**Authors:** Anupama Jha, Mathieu Quesnel-Vallières, Andrei Thomas-Tikhonenko, Kristen W. Lynch, Yoseph Barash

## Abstract

Cancer is a set of diseases characterized by unchecked cell proliferation and invasion of surrounding tissues. The many genes that have been genetically associated with cancer or shown to directly contribute to oncogenesis vary widely between tumor types, but common gene signatures that relate to core cancer pathways have also been identified, signifying that cancer cases display common hallmark molecular features. It is not clear however whether there exist additional sets of genes or transcriptomic features that are less well known in cancer biology but that are also commonly deregulated across several cancer types. Here, in order to agnostically identify transcriptomic features that are commonly shared between cancer types, we used RNA-Seq datasets encompassing thousands of samples from 19 healthy tissue types and 18 solid tumor types to train three feed-forward neural networks, based either on protein-coding gene expression, lncRNA expression or splice junction use, to distinguish between healthy and tumor samples. All three models achieve high precision, recall and accuracy on test sets derived from 13 datasets used during training and on an independent test dataset, indicating that our models recognize transcriptome signatures that are consistent across tumors. Analysis of attribution values extracted from our models reveals that genes that are commonly altered in cancer by expression or splicing variations are under strong evolutionary and selective constraints, suggesting that they have important cellular functions. Importantly, we found that genes composing our cancer transcriptome signatures are not frequently affected by mutations or genomic alterations and that their functions differ widely from the genes genetically associated with cancer. Finally, our results also highlighted that deregulation of RNA-processing genes and aberrant splicing are pervasive features across a large array of solid tumor types. The transcriptomic features that we highlight here define cancer signatures that may reflect causal variations or consequences of disease state, or a combination of both.

## Introduction

Cancer is a loosely defined term that designates cells that have acquired pathological properties, mainly loss of cell cycle regulation, high proliferation rate and loss of contact inhibition leading to invasion of surrounding tissues. In time, tumor cells disrupt the normal function of tissues where they are located and can metastasize to other tissues. Oncogenes contribute to cell transformation while tumor suppressor genes stop aberrant cell proliferation. Changes in expression, activation or function of these genes are expected to lead to cancer-like behavior in various cell or tissue types and many such genes are commonly affected by genomic lesions in cancer. In addition to hallmark oncogenes and tumor suppressor genes, cancer driver mutations that contribute to disease onset and progression are found in subsets of cancer types (Haigis et al., 2019). While these genetic alterations are diverse, several genes that are altered in cancer converge on few molecular mechanisms that are commonly involved in tumorigenesis (Sanchez-Vega et al., 2018). These pathways have wide-ranging effects that span the cell cycle, inflammation and apoptosis, among others. The mechanisms through which they operate in cancer are therefore highly diverse and molecularly heterogeneous, but they are also interrelated. In addition, a recent gene network analysis identified a relatively small number of regulatory modules on which a majority of somatic mutations in cancer converge (Paull et al., 2021). Because changes in cellular pathways and biological activity ultimately impact gene expression and post-transcriptional regulation, this leaves the possibility that tumors that arise from the disruption of different pathways share common molecular signatures in the form of transcriptomic variations.

Previous studies have attempted to leverage these projected common signatures of cancer to train computational models to distinguish tumor from healthy samples or distinguish different tumor types. Typically, these studies rely on protein-coding gene expression data combined with deep neural networks or other machine learning algorithms that classify samples in two or more categories (Dave et al., 2004; Roessler et al., 2010; Wei et al., 2014; Ahn et al., 2018; Grewal et al., 2019; Frost et al., 2019). While these studies show that machine learning models can successfully distinguish between healthy tissues and tumors given a certain set of conditions, the genes used to build these networks are pre-selected to lower the number of biological features used as input and make the training of deep neural network models easier. Although this pre-selection facilitates the training of machine learning models (Blum & Langley, 1997; Guyon & Elisseeff, 2003; Liu & Motoda, 2012), preselecting genes with known functions in the literature or genes already known to be differentially expressed in cancer also deprives the models from learning about potentially novel genes contributing to the transcriptomic signature of cancer.

In addition, while previous studies focused their analyses on the identification of tumor markers, recent methods for the interpretation of deep neural networks offer the opportunity to agnostically discover transcriptomic variations characterizing cancer biology from models that successfully predict biological classes. In particular, we recently described an enhanced integrated gradients method for deep neural network interpretation (Jha et al., 2020) that generates attribution values as a measure of the weight or importance of each biological input feature in the model. For example, we recently used enhanced integrated gradients to find splicing events that are differentially included in brain compared to other tissues without prior knowledge of splicing variations (Jha et al., 2020).

Here, we aimed to draft a molecular profile of cancer that applies to most solid tumor types by leveraging the predictive power of deep neural networks along with the interpretation capability of enhanced integrated gradients to identify common transcriptomic signatures across a large array of tumor types. We trained feed-forward neural networks with protein-coding gene expression, lncRNA gene expression or splice junction usage data from several healthy tissue and tumor types. We then derive attribution values from these models and establish a list of high-attribution features corresponding to a common signature of cancer, which could be causally involved in cancer or result from oncogenic transformation, or both. Finally, we assess the biological functions of these transcriptomic variations.

## Results

### A feed forward neural network trained with protein-coding gene expression distinguishes between healthy and cancer tissues

We aimed to uncover transcriptomic features that commonly define cancer state. Performing differential gene expression analysis on 11 healthy tissue-tumor pairs from GTEx and TCGA and then looking at the overlap in the genes that are deregulated between these analyses shows that no protein-coding gene is consistently up- or down-regulated (abs(log2FC) *>* 2, adjusted p-value *<* 0.01) in more than 9 cancer types (Figure 1A). In addition, while hallmark oncogenes are expected to be disrupted in many cancer types, we observe in a sampling of 11 oncogenes they are often not significantly differentially expressed in any of the 11 cancer analyzed (e.g. BCR, CTNNB1, DDX6, FUS, KRAS, MDM2, TPR) or only disrupted in certain cancer types (e.g. EGFR, ETV4, JUN, MYC; Figure S1, panel A). Although the apparent inconsistency between the function of oncogenes and their lack of change in expression in many tumor types can be partially explained by alternative mechanisms of activation independent of changes in transcript levels, these results demonstrate that using a simple differential gene expression analysis fails to grasp the complexity and heterogeneity of the transcriptomic variations existing across various cancer types.

**Figure 1.**
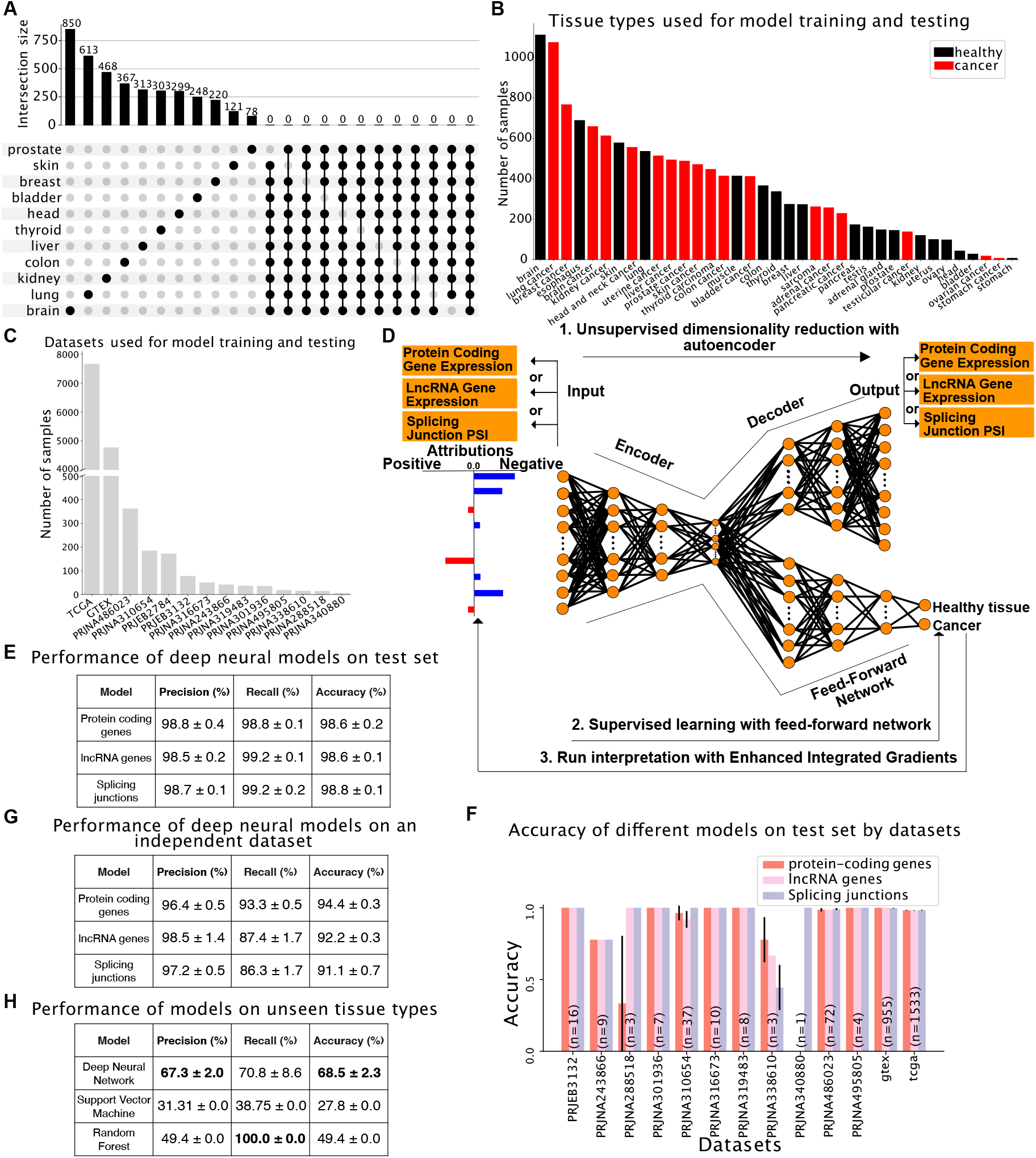
(A) Upset plot summarizing pairwise differential gene expression analyses performed on tumors and their corresponding healthy tissue. No gene is significantly deregulated in more than 9 out of 11 cancer types tested. (B-C) Dataset assembled to train and test binary models to distinguish between healthy and tumor samples shown by tissue type (B) or dataset (C). (D) Graphical representation of the computational framework used to train, test and interpret models. (D) Performance of models trained with protein-coding gene expression, lncRNA gene expression or splicing variations evaluated by precision (number of true positives over the sum of true positives and false positives), recall (number of true positives over the sum of true positives and false negatives) and accuracy (sum of true positives and true negatives over the total population). (E) Performance of models trained with protein-coding gene expression, lncRNA gene expression or splice junction usage on a test set consisting of unseen samples from the 13 datasets used to assemble the training set. (F) Accuracy of models trained with protein-coding gene expression, lncRNA gene expression or splice junction usage across the 13 datasets used to assemble the training set. (G) Performance of models trained with protein-coding gene expression, lncRNA gene expression or splice junction usage on an independent dataset consisting of healthy and cancer lung samples. (H) Performance of the deep learning model, SVM and Random Forest using protein-coding gene expression on unseen tissue types with no batch correction and liquid tumors. The training set consists of solid tumors only.

In order to overcome these limitations and identify what transcriptomic features are important in the determination of cancer state, we sought to train interpretable deep learning models capable of distinguishing between healthy and cancer samples. We assembled a large RNA-Seq dataset comprising 13461 samples from 19 healthy tissue types and 18 cancer types (Figure 1B). Samples were sourced from TCGA (https://www.cancer.gov/tcga), GTEx (Lonsdale et al., 2013) and 12 other datasets (Figure 1C). Because technical biases and batch effects are a major concern when using large-scale RNA-Seq datasets, especially when comparing perfectly confounded datasets like GTEx and TCGA, we included in our compendium 12 smaller datasets containing either only tumor samples or tumor and matched healthy tissue samples from the same donors. These additional datasets allow us to mitigate dataset-specific biases and focus on cancer-specific signals by performing a tissue/tumor-specific mean correction across the 14 datasets (see Figure S1, panel B).

We first used expression data from 19657 protein-coding genes to train an autoencoder for dimensionality reduction, followed by a supervised neural network to predict cancer state (healthy tissue versus tumor, Figure 1D; see Methods for details about dimensionality reduction and model training). To ensure that our model did not learn dataset-specific biases, we evaluated model performance on a previously unseen set of samples extracted from the 13 datasets used for training as well as one independent dataset (PRJEB2784) that was not used during training but that comprises 172 samples of tissue types that were included in the training set (healthy and tumor lung samples). Our protein-coding gene expression model accurately predicted whether an individual sample corresponds to a healthy tissue or a tumor (accuracy: 98.6% ± 0.2%, precision: 98.8% ± 0.4% and recall: 98.8% ± 0.1%, Figure 1E) and this performance generalizes over all 13 datasets despite the class imbalance between datasets (Figure 1F, see numbers in the bars for the number of samples in each dataset). The only datasets where the performance is variable have very few examples (PRJNA340880 has 1 sample and PRJNA288518 has 3 samples). Importantly, the model performs almost as well when applied on the independent dataset (Figure 1G).

To evaluate how our model generalizes to cancer types not included in the training set, we assembled a group of healthy (T and B lymphocytes) and malignant (acute myeloid leukemia and acute lymphoblastic leukemia) blood cells from three datasets (ArrayExpress E-MTAB-2319, Blueprint and TARGET consortia, see Supplementary Table 1 for details on the samples). We omitted batch correction on these samples to assess how our model would fare on a dataset that considerably differs from the training set both at the biological (solid tissues and tumors in training set vs. hematologic cells and tumors in test set) and technical levels (batch-corrected in training set vs. uncorrected test set). Strikingly, despite these vast differences between training and test sets, our deep neural network model successfully distinguishes healthy and cancer samples from blood (Figure 1H), although at lower accuracy, as expected due to the biological and technical differences between the training and test sets.

Finally, in order to assess how our deep neural network model compares to other machine learning algorithms, we first trained Support-Vector Machine and Random Forest models using the same training set as for our deep neutral network model, and then tested them on the same independent dataset consisting of batch-corrected healthy and cancer lung samples. Under these conditions, all three models perform similarly (Supplementary Table 7A). However, in sharp contrast with the deep neural network model, both the Support-Vector Machine and Random Forest models completely fail when applied to the hematologic dataset (Figure 1H). These results demonstrate that a deep neural network trained on protein-coding gene expression profiles can robustly predict whether a sample comes from a healthy tissue or a tumor, even when tested with tissue types not previously seen, and that deep neural networks perform better than other algorithms for this task.

### lncRNA expression or splice site usage profiles suffice to define cancer state

Other types of transcriptomic features, including lncRNA expression and RNA splicing, have been used as prognostic markers or to predict drug response in cancer (Ching et al., 2016; Bolha et al., 2017; Brinkman, 2004). In addition, a small number of mutations located in lncRNA genes or disrupting splicing in protein-coding genes have been shown to drive cancer (Carlevaro-Fita et al., 2020). However, it is not known whether widespread changes in lncRNA expression or RNA splicing commonly characterize cancer state. We thus asked if these other types of transcriptomic features could be used to distinguish between healthy and tumor samples, similarly to what we found for protein-coding gene expression.

We used the same strategy as for the model trained with protein-coding gene expression above and trained models with expression data from 14257 lncRNA genes or splice site usage data from 40147 splice junctions. Remarkably, these models achieved 98.6% ± 0.2% and 98.8% ± 0.1% accuracy, respectively, with high precision (98.5% ± 0.2% for lncRNA expression and 98.7% ± 0.1% for splice junction usage) and recall (99.2% ± 0.1% for lncRNA expression and 99.2% ± 0.2% for splice junction usage, Figure 1E). As observed with the protein-coding gene expression-trained model, the lncRNA gene expression and the splice junction usage-trained models perform consistently well across all of the test datasets on the task of predicting the cancer state, again despite the class imbalance (Figure 1F), or on an independent dataset (Figure 1G), showing that our models predict classes based on biological signal and not confounding factors.

As for the protein coding gene model, we compared our deep neural network model for lncRNA gene expression with Support-Vector machine and Random Forest models on the same two datasets. All three models performed well on the batch-corrected dataset consisting of healthy and tumor lung samples (Supplementary Table 7B). However, while our deep neural network model also correctly predicted cancer state with the hematologic dataset (healthy lymphocytes and leukemia samples), both Support-Vector machine and Random Forest completely failed again on this dataset (Supplementary Table 8). On the splicing data, our deep neural network model outperforms both the Support-Vector machine and Random forest models on the independent batch-corrected healthy and tumor lung dataset (Supplementary Table 7C). The strong performance of our lncRNA- and splicing-trained models indicates that tumor samples can be defined not only by their protein-coding gene expression profile, but also using exclusively their lncRNA gene expression or splice junction usage profile.

### Interpretation of deep learning networks uncovers new transcriptomic features characterizing cancer state

Given the high performance of our models, we wanted to know what transcriptomic features are the most important in each of our models and whether these features consist mostly of the usual suspects, i.e. genes known to be genetically associated with cancer. To do this, we generated feature importance scores known as attribution values for tumor samples using enhanced integrated gradients (EIG) (Jha et al., 2020). Briefly, EIG measures a feature’s contribution, either positive or negative, to the model label predictions (healthy tissue versus cancer) when comparing a cancer sample to a baseline. Following our previous work (Jha et al., 2020), we used the median of normal samples as the baseline (see Methods section “Interpretation of tumor classification models” for details).

We selected 1768 protein-coding genes, 1763 lncRNAs and 562 splice junctions that have high median attribution values across tumor types (Figure 2A; see Methods section “Selection of feature sets” for selection criteria and Figure S2). We also defined “neutral” sets with a sample size equivalent to sets of high-attribution features using features that display attribution values close to zero. When looking at the cancer type-specific attribution values across 14 tumor types for the top 100 features with positive or negative attribution, we found that protein-coding genes, lncRNAs and splice junctions with the highest median attribution values across all tumor samples have consistently high attribution values in most if not all cancer types (Figure 2B), highlighting that our models are not driven by outlier expression or splice junctions usage in cancer types with a large sample size, but rather rely on common transcriptomic features of cancer.

**Figure 2.**
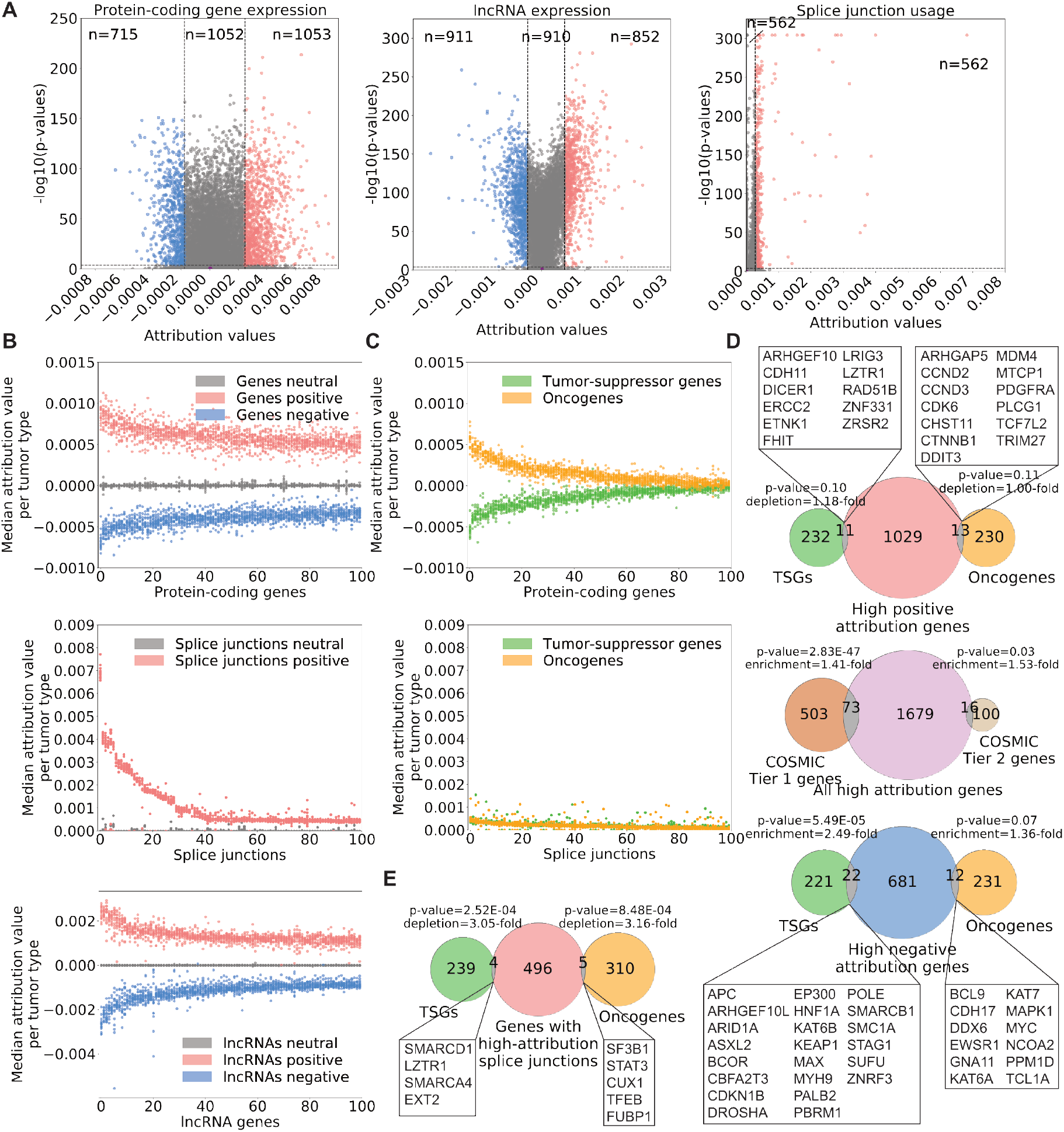
(A) Selection of high-attribution features from models trained with protein-coding gene expression, lncRNA gene expression or splice junction usage. Dotted lines show cutoffs used; purple points around coordinates (0,0) show features selected in neutral sets. (B) Median attribution values of 100 protein-coding genes, lncRNAs or splice junctions with the highest positive and negative attributions across tumor tissues. (C) Median attribution values of genes associated with cancer from the COSMIC database. (D) Overlap between COSMIC oncogenes and TSGs, and genes with high positive (top panel) or negative (bottom panel) attribution values, or between COSMIC tier 1 genes (high confidence for causal role in cancer) and tier 2 genes (some evidence of causal involvement in cancer), and all high-attribution genes (central panel). (E) Overlap between genes associated with cancer and genes harboring junctions with high attribution values. In both D and E, enrichment or depletion factors were calculated from the ratio of observed vs. expected overlapping genes between sets, and p-values were calculated from hypergeometric tests.

In agreement with our differential gene expression analysis that showed that no gene is significantly deregulated in the same manner across all tumor types, we find that the sign of the attribution values does not necessarily reflect the change in expression in cancer. In other words, a gene with a high positive attribution value won’t necessarily be upregulated in all or most cancers, and conversely, a gene with a high negative attribution value won’t necessarily be downregulated in all or most cancers. Thus, rather than highlighting genes and splicing variations that are similarly altered in many cancer types, the interpretation of our models exposes transcriptomic variations that consistently deviate from the norm in cancer. Since the models are not causal, the transcriptomic variations that they identify could reflect changes that cause or are a consequence of cancer, or a mixture of both.

We next sought to assess the relation between model attributions and known oncogenes or tumor suppressor genes (TSGs). Strikingly, we found a clear separation between the latter two groups with oncogenes receiving positive attribution and TSGs receiving negative values (Figure 2C). However, most the known oncogenes and TSGs have moderate attribution values relative to our top-scoring features, with many having neutral attribution values (close to 0). This result was observed for both the expression of COSMIC genes or usage of splice junctions found of those genes. Compared to our criteria for selection of high-attribution features we only observe a small, although statistically significant, enrichment for COSMIC genes among our high negative attribution genes (Figure 2D). Of note, well-known oncogenes and TSGs are depleted among genes that have splice junctions with high attributions, meaning that there are fewer oncogenes and TSGs with high-attribution splice junctions than would be expected by chance (Figure 2E). These results show that our models rely on gene expression and splicing variations in genes that mostly differ from established oncogenes and TSGs to predict tumor samples, and that the transcriptomic definition of cancer that we provide here largely differs from genes harboring hallmark mutations causally implicated in cancer.

### High evolutionary and selective constraints in transcriptomic features defining tumor state

After establishing a list of genes with high attribution values by expression or splice junction usage and discovering that most of these genes do not correspond to COSMIC oncogenes or TSGs, we wondered whether transcriptomic features that carry high attribution values in our models have properties that may indicate important roles in cells. We discovered that protein-coding genes, lncRNA genes and genes with splice junctions corresponding to high-attribution features in our models are highly evolutionarily conserved relative to the neutral sets (Figure 3A). We noted that proteincoding genes that have high negative attributions as well as lncRNA genes that have high positive or negative attributions are in general significantly longer than the reference neutral sets, but that genes with splice junctions with high attributions are significantly shorter (Figure 3B). We also observed that protein-coding genes and genes with splice junctions with high attributions display high selective pressure against loss of function mutations, as estimated by the gnomAD loss-of-function observed/expected upper bound fraction (LOEUF) score (Karczewski et al., 2020) (Figure 3C).

**Figure 3.**
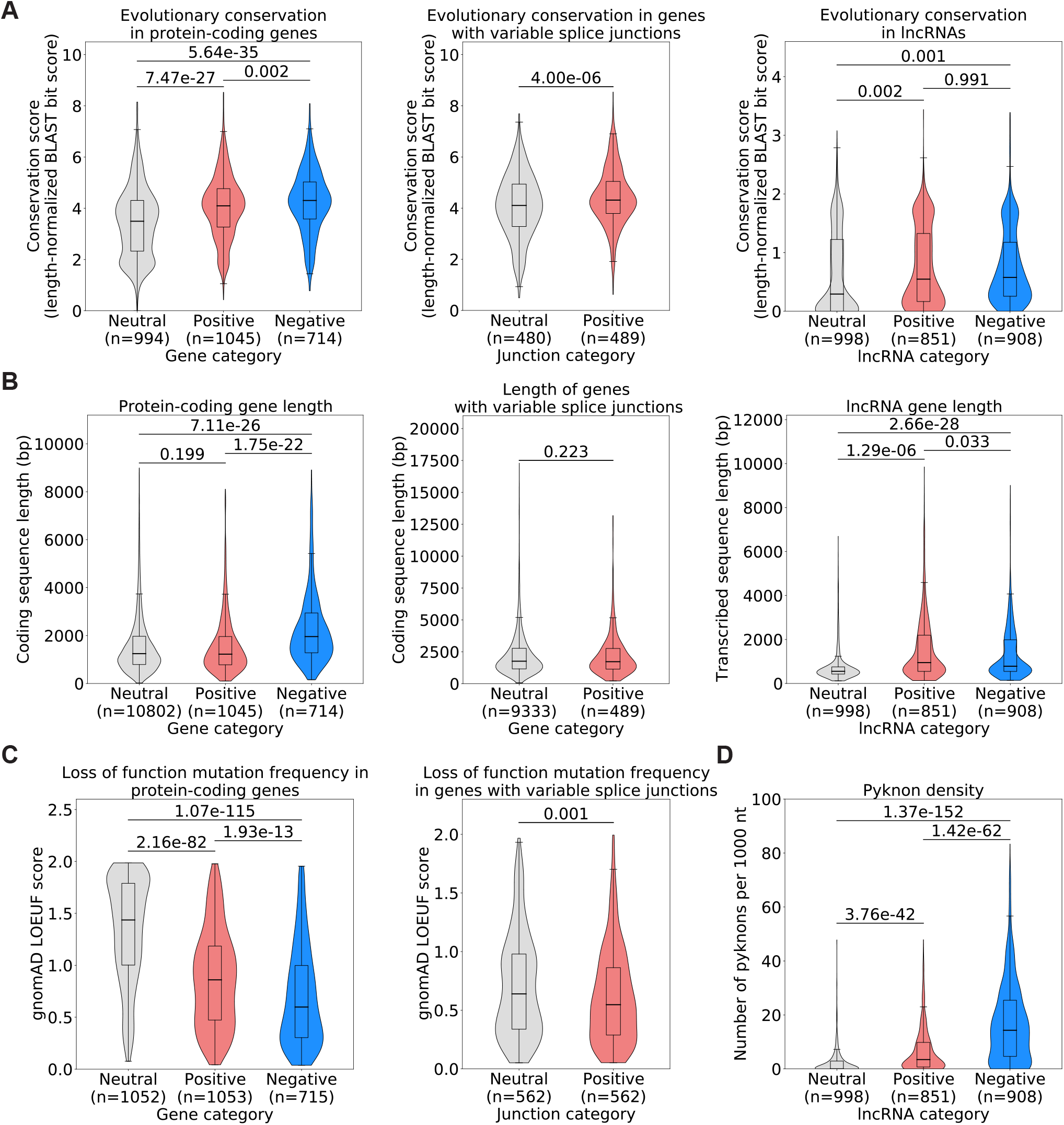
(A) Evolutionary conservation of protein-coding genes (left panel), genes with variable splice junctions (middle panel) or lncRNA genes (right panel) in human, chimpanzee, mouse, cattle, xenopus, zebrafish and chicken. (B) Gene length derived from the longest annotated transcript in Ensembl (hg38) for protein-coding genes (left panel), genes with variable splice junctions (middle panel) or lncRNA genes (right panel). (C) Selective pressure against loss-of-function mutations in the human population as assessed by gnomAD LOEUF score (Karczewski et al., 2020), which corresponds to the frequency at which loss-of-function mutations are observed in the human population for a given gene, showing score for protein-coding genes (left panel) or genes with splice junctions whose usage we used for model training (right panel). A low LOEUF score means high selective pressure against loss of function. (D) Pyknon density in lncRNA genes whose expression we used for model training. All statistics: unpaired t-test.

Finally, we inferred the functional impact of lncRNA genes with high attributions by looking at their density of a class of DNA motifs termed pyknons. Pyknons are located in loci that were previously reported as often differentially transcribed between healthy and colorectal cancer tissues and that can affect the oncogenic functions of lncRNAs (Rigoutsos et al., 2006; 2017; Evangelista et al., 2020). We found that high-attribution lncRNA genes carry a higher density of pyknons (Figure 3D) than lncRNA genes from the neutral set. This was true for both positive- and negative-attribution lncRNAs, but it was particularly marked in negative-attribution lncRNAs, where average pyknon density is seven times higher than neutral-attribution lncRNAs. Together, these findings show that high attribution proteincoding and lncRNA genes by expression or splicing are under strong evolutionary and selective constraints and suggest that these protein-coding genes and lncRNAs with altered expression or abnormal splicing in cancer have essential functions in cells.

### Characterization of splice junctions with high attributions

While it is easy to conceive how changes in the expression level of a gene can drive tumorigenesis, interpreting the impact of splicing changes in disease is not as straightforward. We thus wanted to assess how variable splice junctions with high attributions are predicted to impact protein sequence and function. We first noted that high-attribution junctions are predicted to disrupt the reading frame of the gene as often as our reference neutral junction set (Figure S3, panel A). We also looked at predicted disorderness of the peptide sequence corresponding to the two exons immediately upstream and downstream of variable splice junctions and found that the predicted peptide disorderness level is no different in high-attribution junctions from what we observe in the neutral set (Figure S3, panel B).

We then verified whether high-attribution splice junctions affect known protein domains by predicting the protein domains encoded from the two exons immediately upstream and downstream of high-attribution junctions using the NCBI Converved Domain Database. Interestingly, we discovered that 11 splice junctions in 10 genes (*CSNK2A2, MAPK9, RIOK1, PRKDC, TYK2, PAK1, IRAK1, CSNK2A1, VRK1, MARK3*) affect a part of the transcript matching sequences of protein kinase C (PKC)-like superfamily domains (Figure 4). While genes contributing to PKC signaling have been implicated in cancer as oncogenes or tumor suppressors (Garg et al., 2013), little is known about the impact of splicing variations altering PKC-like superfamily domains in cancer. We also found additional high-attribution splice junctions that affect other domains that are linked to cancer signaling, in particular DEAD-like, RING and C2 domains. Thus, it is possible that some of the high-attribution splice junctions that we uncovered regulate cancer through the alterations of cancer signaling protein domains.

**Figure 4.**
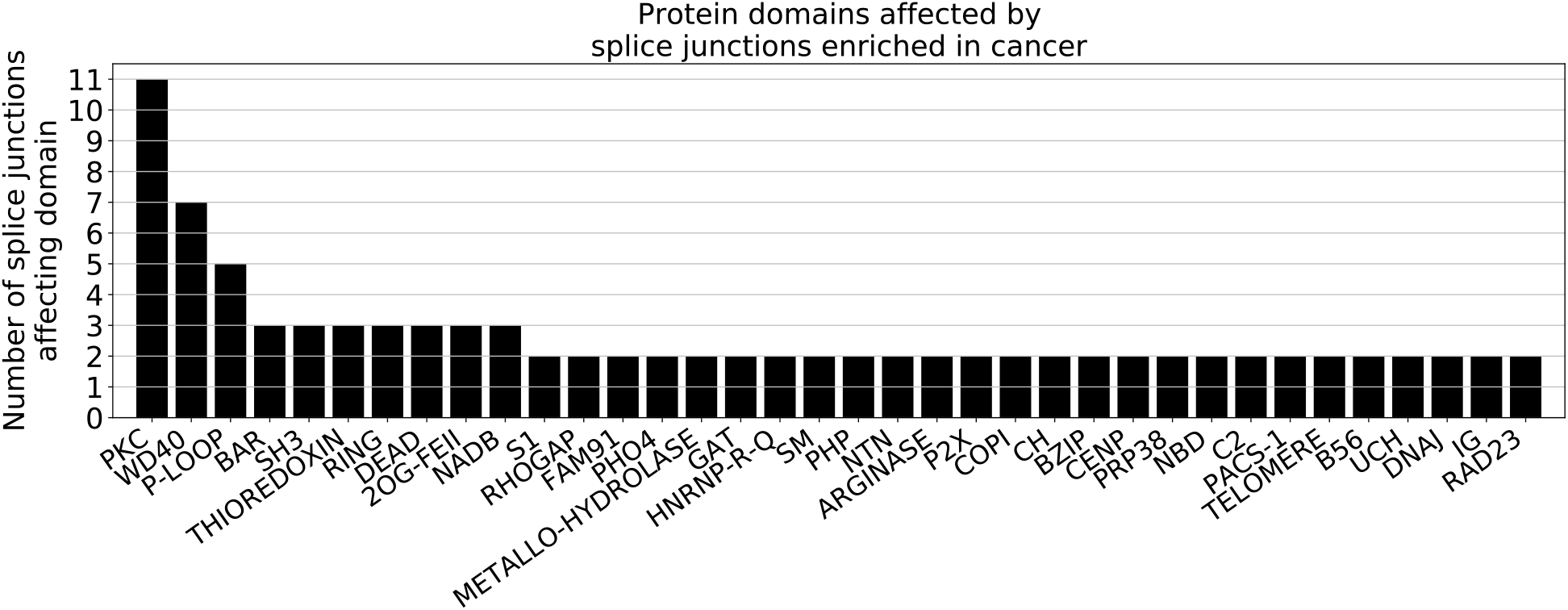
Protein domains that are affected by at least two splice junctions with high attributions.)

### Frequency of genetic alterations in transcriptomic features characterizing cancer state

Next, we wondered if genetic alterations in our high-attribution genes might be driving the transcriptomic variations highlighted by our models. We first postulated that high-attribution genes would rarely carry driver mutations since these genes are not known to be genetically linked to cancer, which we confirmed by looking at TCGA samples and finding that less than 2% of high-attribution genes carry a driver mutation in at least one of any of the samples in TCGA (Figure 5A). While high-attribution genes do not carry driver mutations, our analysis shows that genes with high negative attribution values by expression display a higher frequency of passenger mutations than their reference neutral set, and that the frequency of passenger mutations in high negative attribution genes is as high as in COSMIC oncogenes (Figure 5B). The frequency of structural variants, although higher in high-attribution genes than their reference neutral sets, is lower for all sets of high-attribution genes than for COSMIC genes (Figure 5C). Similarly, the frequency at which high-attribution genes are involved in amplification (Figure 5D) or deletion events (Figure 5E) is not significantly different from the neutral sets or the COSMIC genes. Overall we conclude that the cancer transcriptomic features we identified are not frequently affected by genetic alterations, which suggests that the cancer expression and splicing patterns obtained from our models are not driven by genetic variations in these genes.

**Figure 5.**
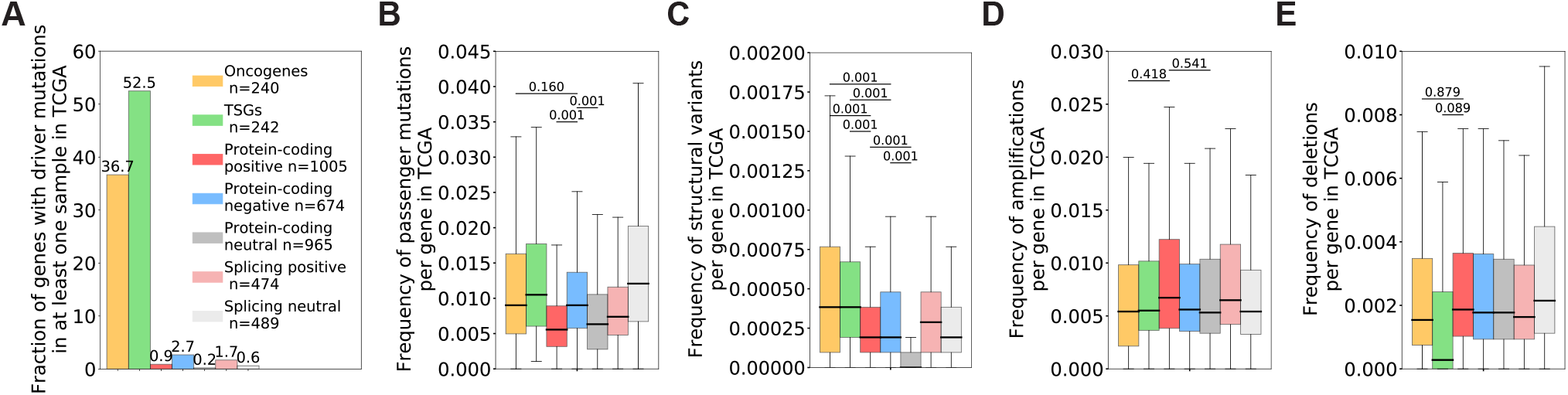
Fraction of genes with driver mutations (A), or frequency of passenger mutations (B), structural variants (C), amplification events (D) or deletion events (E) in the TCGA cohort for COSMIC oncogenes, COSMIC tumor-suppressor genes (TSGs), genes with high positive (“Protein-coding positive”), high negative (“Protein-coding negative”) or neutral attribution values by expression (“Protein-coding neutral”), or genes with splice junctions with high (“Splicing high”) or neutral (“Splicing neutral) attribution values in our models.One-way ANOVA with Tukey post-hoc tests.

### Contrasting functions of genes with high positive or negative attributions by expression or splicing in cancer

Finally, given that a majority of the protein-coding genes or genes with splice junctions with high attribution values in our models were not previously associated with cancer, we sought to understand the functions of those genes. We first checked whether genes that have high attribution values by expression differ from the genes that have high attributions by splice junction usage and confirmed that a large majority of the genes with high attributions by expression differ from the genes with high attributions by splice junction usage (Figure 6A). We performed a gene ontology analysis for protein-coding genes with high attribution values and found that protein-coding genes with high negative attribution values are enriched for functions related to transcription, mitosis, histone modification, chromatin regulation, and localization to the centrosome, in line with the traditional view of cancer (Figure S4, panel A). In sharp contrast, protein-coding genes with high positive attribution values are enriched for post-transcriptional and post-translational modifications, in particular tRNA modification, RNA splicing and protein neddylation, as well as membrane-bound organelles (Figure S4, panel B). Similar to protein-coding genes with high positive attribution values, genes with splice junctions with high attribution values are also enriched for functions related to RNA processing, in particular splicing and transport, but also carry a component of terms related to chromatin (Figure S4, panel C). Of note, the set of genes with neutral attribution values by expression are enriched for heterogeneous and unrelated terms (Figure S4, panel D) and genes with neutral attribution values by splice junction usage failed to return any terms with an adjusted p-value *<* 0.05, indicating that our high-attribution genes consist of sets of biologically related and consistent functions.

**Figure 6.**
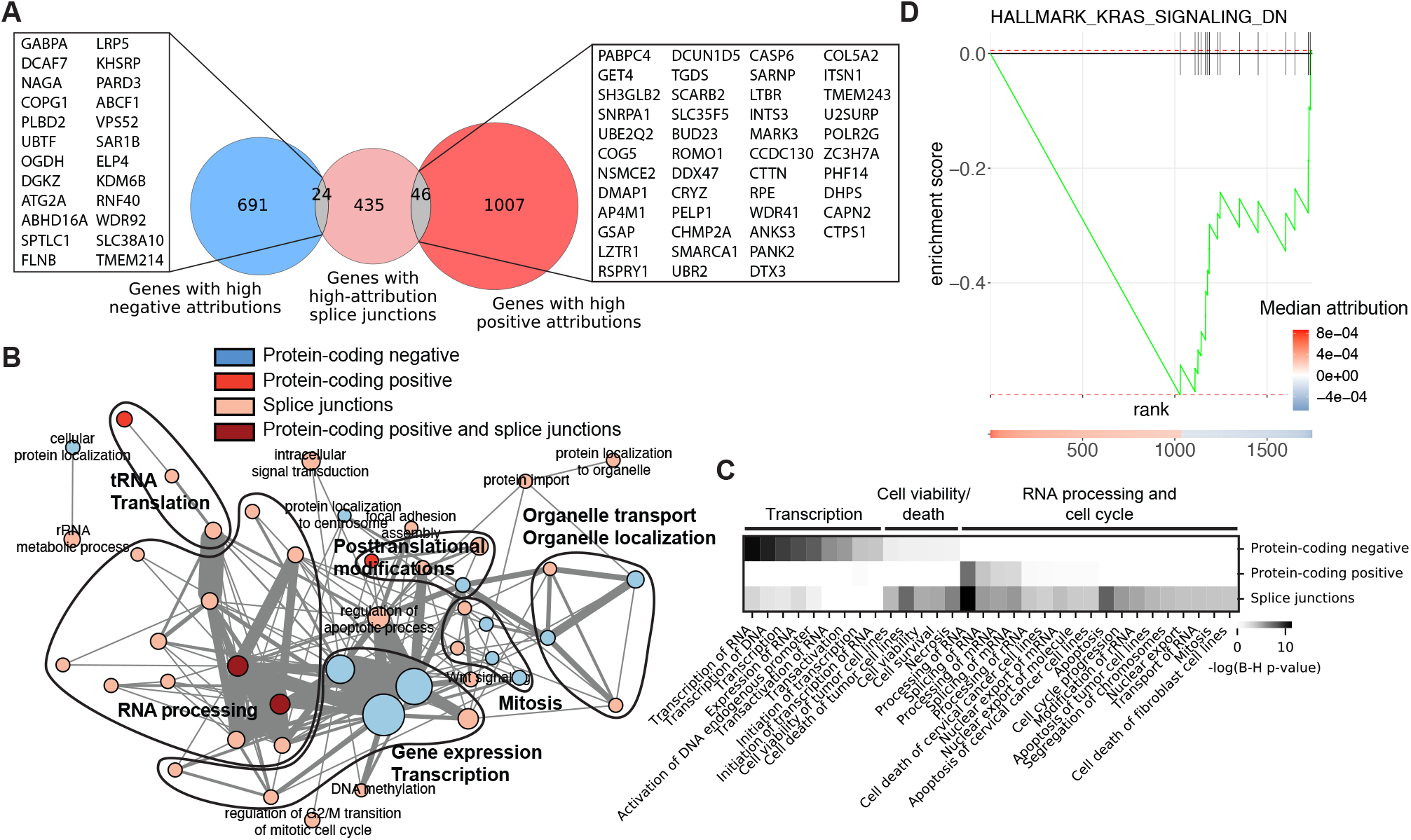
(A) Overlap between genes with high positive (red set) or negative (blue set) attribution values and genes with splice junctions that have high attribution values (light red set) with the list of genes overlapping between sets. (B) Enrichment map showing GO terms related to biological processes that are enriched among protein-coding genes with high negative (Protein-coding negative, blue) or positive (protein-coding positive, red) attributions by expression, genes with high-attribution splice junctions (splice junctions, light red), or enriched both in protein-coding positive and splice junction genes (protein-coding positive and splice junctions, dark red). Each node is a GO term and the color of the nodes corresponds to gene sets in which they are enriched. The thickness of edges corresponds to the number of genes in common between GO terms. (C) Heatmap of terms obtained from an Ingenuity Pathway Analysis (QIAGEN) for molecular and cellular functions. (D) Gene set enrichment analysis on high-attribution protein-coding genes by expression showing an enrichment in KRAS signaling among negative attribution genes. High-attribution genes were ranked on the x-axis from high positive (left, red) to high negative attribution (right, blue).

An enrichment map built from GO terms related to biological processes shows that high positive attribution genes by expression or splicing form a highly interconnected network whose core relates to functions associated with RNA biology (Figure 6B). The core network from high positive attribution genes by expression or splicing differs from the highly interconnected network core drawn from genes with high negative attributions by expression, which are associated with gene expression and transcription (Figure 6B). Ingenuity Pathway Analysis for molecular and cellular functions associated with high-attribution genes confirmed that functions of high negative attribution genes are often distinct from those of high positive attribution genes, with transcription and RNA processing being predominant, respectively (Figure 6C). Pathway analysis also revealed how high negative attribution genes by expression and genes with high attribution by splice junction usage overlap in functions related to cell survival and how high attribution genes by splice junction usage are involved in the cell cycle. Finally, gene set enrichment analysis unveiled an enrichment of high negative attribution genes for KRAS signaling (Figure 6D), while no significant enrichment was found for genes with high positive attributions by expression or splicing.

Thus, while genes that have high negative attribution values in cancer share functions of known oncogenes and TSGs, including in how they are implicated in genome maintenance and transcription, genes that have high positive attributions by expression or splicing have distinct functions, several of which are related to RNA regulation and RNA processing.

## Discussion

Our results demonstrate that feed-forward neural networks can be used to distinguish between healthy and tumor samples using transcriptomic features. Importantly, we show that models trained with lncRNA expression or splice junction usage perform as well as, if not better than, a model trained with protein-coding expression data. This observation highlights how various elements of the transcriptome can inform on disease state and emphasizes the importance of pursuing molecular markers beyond variations in proteincoding gene expression, in particular by assessing variations in lncRNA gene expression and splice junction usage in cancer. Our approach uncovered common transcriptomic profiles consisting of a number of gene expression and splicing variation markers that are not altered in the same way across all solid tumor types, making up a novel molecular definition of cancer that would be impossible to establish using traditional approaches such as differential gene expression analysis.

The interpretation of our models revealed known and novel molecular features of cancer. Known cancer drivers were enriched among genes that we find to have high attribution values, which can be expected for genes with driver mutations resulting in loss of function or lower protein expression. In addition, genes with high negative attribution values comprised genes with functions typically associated with genome integrity maintenance, such as histone modification and chromatin regulation, as well as transcription, two long-known aspects of cancer development (Hanahan & Weinberg, 2000; 2011). On the other hand, many of the protein-coding genes that have high positive attribution values have roles in RNA regulation or RNA processing. Interestingly, RNA deregulation has become a recurrent theme in cancer research (Goodall & Wickramasinghe, 2020). Driver mutations have been found in several RNA-binding proteins (e.g. *SF3B1, U2AF1, SRSF2, HNRNPA2B1, SRRM2*) in cancers ranging from blood malignancies to glioblastoma (Anczuków & Krainer, 2016; Marabti & Younis, 2018; Sveen et al., 2015), but several questions remain regarding how widely this group of proteins and their targets are involved in cancer. Our results suggest that RNA deregulation might be a central component of cancer, upon which many cellular pathways involved in cancer may converge. Indeed, network analysis shows that genes with high attribution values by expression or splice junction usage and that have functions related to RNA regulation are tightly connected to the canonical pathways of cancer (Figure 7, Supplementary Table 9).

**Figure 7.**
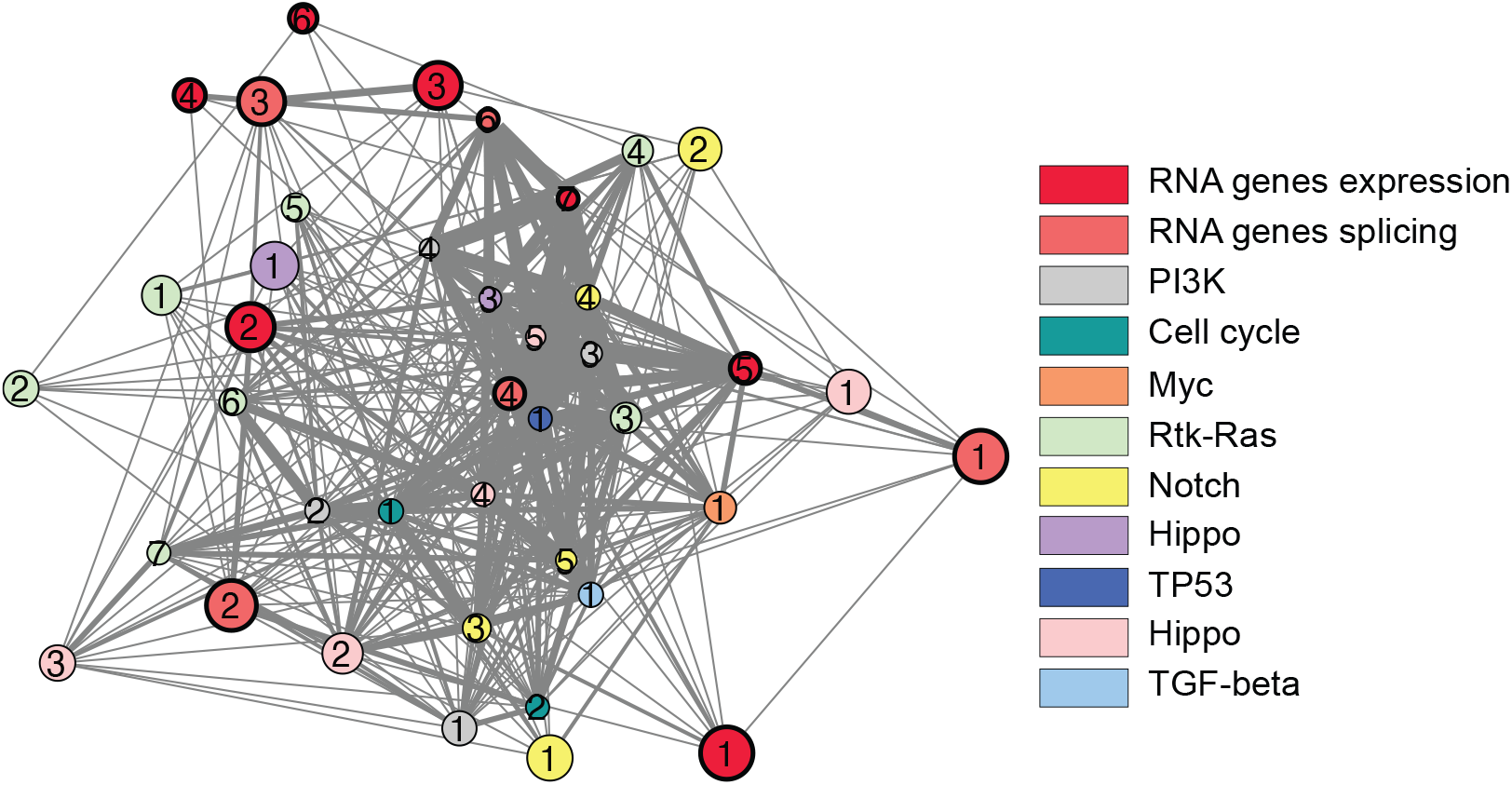
Network analysis of common cancer pathways (PI3K, cell cycle, Myc, Rtk-Ras, Notch, Hippo, TP53, Hippo, TGF-beta) together with genes in GO terms related to RNA regulation or RNA processing that are enriched in our protein-coding or splicing models. Each node is a network as predicted with Ingenuity Pathway Analysis; the size of the nodes represents the number of molecules in each network and the thickness of the edges represents the number of molecules in common between two networks. Networks formed by high-attribution genes in our protein-coding and splicing models are highlighted with a thicker node border. Only networks comprising at least three molecules and connected by at least two shared molecules are shown. Numbers identify networks; Supplementary Table 9 lists the molecules found in each network (node).

Our transcriptomic definition of cancer includes several elements that were not previously genetically associated with cancer but that display strong sequence constraints, which suggests that these genes or splice junctions play essential roles in cells. Interestingly, several of these genes that are not listed as COSMIC oncogenes still display tumorigenic characteristics. A few examples include *DYNC1H1* (Gong et al., 2019), *WSB1* (Kim et al., 2016; Cao et al., 2015; Kim et al., 2015), *RUFY3* (Xie et al., 2017; Wang et al., 2015), *DOCK5* (Frank et al., 2016), *MYSM1* (Li et al., 2017), *DSE* (Liao et al., 2018), *DCUN1D5* (Bommeljé et al., 2013; Guo et al., 2012), *SARNP* (Kang et al., 2019) and *FNTA* (Tian et al., 2019; Chen et al., 2020a), which can all promote cell proliferation or transformation, at least in some conditions. Likewise, we have identified splice junctions in cancer that deviate from normal tissues in functional domains implicated in cancer signaling, such as PKC-like (Garg et al., 2013), DEAD-like (Fuller-Pace, 2013) and RING (Lipkowitz & Weissman, 2011) domains, in genes associated with cancer, such as *CSNK2A1, CSNK2A2, RIOK1, PRKDC, TYK2, PAK1* and *IRAK1*, for which gene expression and posttranslational modifications act as mechanisms for cancer progression (Zhang et al., 2019; Hong et al., 2018; Gray et al., 2014; Wöss et al., 2019; Ye & Field, 2012; Cheng et al., 2018). However, splicing variations related to cancer have not been reported for any of these genes except *PAK1*, for which a JMJD6-regulated exon inclusion altering the PKC domain enhances MAPK signaling in melanoma (Liu et al., 2017). While the splice junction we identified differs from the one reported before, it also affect the PKC domain of PAK1. *IRAK1* has two well-characterized alternative splicing events (Jain et al., 2014), but there exists no evidence that these events are directly involved in tumorigenesis, and they also differ from our splice junctions with high attributions.

Interestingly, while our analysis of variant frequency shows that high negative attribution genes in our protein-coding gene expression model are more frequently mutated than the neutral set, and almost as frequently as COSMIC oncogenes and TSGs, variant frequency is much lower for genes with high positive attributions by expression or splicing, which could explain why many of these transcriptomic features have previously been overlooked in genomic studies.

Future work should be directed at assessing whether these and additional novel transcriptomic features of cancer that we highlight here are causally involved in tumorigenesis, thereby highlighting a core transcriptomic component of cancer development, or whether they represent the downstream consequences of genetic alterations driving cancer, and how these variations relate to core transcriptional circuitries of cancer (Chen et al., 2020b).

## Methods

### Notation

We are interested in defining the transcriptomic signature of solid tumors. We achieve this by first predicting the cancer state of an RNA-seq sample using a deep learning model with gene expression (protein-coding or lncRNA) or splicing quantification as input. Subsequently we interpret the prediction made by the deep learning model given our input observation by assigning attributions to each feature of the observation. Let 𝒳 be the input space and 𝒴 be the output or label space. Input ***x*** is in a *p*-dimensional feature space 𝒳 = ℝ^*p*^. Since we only consider a binary classification task for defining the cancer state, 𝒴 ={0, 1}, where 0 represents healthy tissue and 1 represents tumor. Predictions are obtained by a prediction function on the feature space *F* : 𝒳 → 𝒴. The goal of the interpretation step is to obtain a *p*-dimensional vector of attributions called **attr** ∈ ℝ^*p*^, with each value representing how each of the *p* features contributes to the prediction *F* (***x***).

### Datasets

In this work, we process RNA-seq samples from healthy human tissues and tumors from 14 datasets (see Supplementary Table 1 for a list of all samples and their tissue/cancer identity and figure 1A and 1B for the number of samples in each dataset and their cancer state). The two largest datasets among these are from the Genotype-Tissue Expression (GTEx) consortium and the Cancer Genome Atlas (TCGA) program. We processed 7665 samples from 19 healthy tissues from GTEx and 4777 tumor samples from 18 cancer types from TCGA. Since large datasets often suffer from batch effects, we included 12 other datasets in our analysis (Ju et al., 2012; Seo et al., 2012; Lee et al., 2016; Kirby et al., 2016; Ooi et al., 2016; Yang et al., 2017; Yun et al., 2017; Lin et al., 2019; Corona et al., 2020; Qin et al., 2020). Two of these datasets included lung tumor samples with matched healthy lung samples, two included liver tumor samples with matched healthy liver samples, one included stomach tumor samples with matched healthy stomach samples, one included breast tumor samples with matched healthy breast samples, one included brain tumor samples with matched healthy brain samples, one included colon tumor samples with matched healthy colon samples, one included only head and neck cancer samples, one included only pancreatic tumor samples, one included only ovarian tumor samples and one included only prostrate tumor samples. For testing on independent, unseen tissue and tumor types, we processed 16 macrophage samples, 8 monocyte samples and 9 CD4^+^ lymphocyte samples from the Blueprint project (Fernández et al., 2016), 35 CD4^+^ and 14 CD8^+^ lymphocyte samples from dataset E-MTAB-2319 (Ranzani et al., 2015), and 40 B-cell acute lymphoblastic leukemia samples and 40 acute myeloid leukemia from the pediatric TARGET cohort (Ma et al., 2018).

### RNA-Seq data processing

In order to minimize the introduction of technical biases, all RNA-Seq samples were processed from fastq files (legacy TCGA) in the same manner. The raw reads from RNA-Seq experiments are passed through quality control using FastQC (Andrews et al., 2010). Sequencing adapters were trimmed with TrimGalore (v0.6.6) (Krueger, 2018), reads were aligned with STAR (v2.5.2a) (Dobin et al., 2013) against the hg38 human genome assembly (Hubbard et al., 2002), and mapped reads were sorted and indexed using samtools (v1.11). Gene expression quantification was carried out using Salmon (v0.14.0) in quasi-mapping mode using an index generated with Ensembl GRCh38 transcriptome release 94. Gene expression estimates (transcripts per million) are generated from the STAR output using Salmon (Patro et al., 2017). Transcripts per million (TPM) is a normalization method for RNA-Seq data described as:

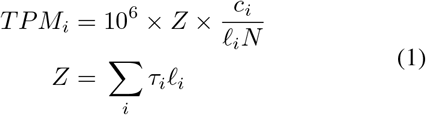

where *TPM*_*i*_ is transcript per million for transcript *i, c*_*i*_ is number of reads mapped to transcript *i, N* is the total number of mappable reads, 𝓁_*i*_ is the length of transcript *i*, 𝒯_*i*_ is the expression level (relative abundance) of transcript *i*, and *Z* is the normalization factor (mean length of expressed transcripts) (Dewey et al., 2019). Most genes have more than one transcript and we summed all of the transcript TPMs into a single TPM value per gene (Love et al., 2017). Splice junctions were quantified using MAJIQ (v2.1) (Vaquero-Garcia et al., 2016). MAJIQ defines alternative splicing in terms of local splicing variations (LSVs). LSVs can be binary or complex. Binary LSVs comprise of only two junctions and complex LSVs have more than two junctions. We only use binary splicing variations that were quantified in at least 80% of samples of a given tissue or tumor type and select junction usage value randomly from one of the two junctions composing the splicing variation. The splicing quantification of a junction in a condition is called percent spliced-in (PSI). For binary LSVs, PSI measures the ratio of the number of reads supporting inclusion of a junction in a condition over the total number of reads supporting its inclusion or exclusion. MAJIQ uses a Beta-binomial distribution over the reads to quantify PSI. For more details on the statistical model please refer to (Vaquero-Garcia et al., 2016).

### Batch correction

To mitigate batch effects from our RNA-seq data, we take two steps. First, in addition to GTEx and TCGA, we searched for other datasets with healthy tissues and tumor samples. This step is necessary as our signal of healthy tissues versus tumor is confounded with whether the sample came from GTEx or TCGA. GTEx consists of healthy tissues and TCGA consists of mostly tumor samples, therefore, we added 12 other small datasets to ensure that we learn cancer-specific signal and are not confounded by dataset-specific technical biases. Next, we correct dataset bias for gene expression data by mean-centering each tissue/tumor-type separately across datasets. Batch correction step ensures our deep learning model using proteincoding and lncRNA gene expression generalizes across multiple datasets.

### Tumor classification models

We train three deep learning models for defining the transcriptomic signature of solid tumor samples. Each of these models use a different set of transcriptomic features to predict the cancer state of an RNA-seq sample: protein-coding gene expression, lncRNA gene expression and quantification of splice junctions. Since we have thousands of noisy transcriptomic features, we first train an autoencoder model for dimensionality reduction followed by a supervised feed forward neural network for the cancer state prediction. In the next three sections, we describe these models in detail.

### Protein-coding gene expression based model

In our first model, the input features are the expression values of 19657 protein coding genes. We extracted the protein-coding genes from Ensembl BioMart (see Supplementary Table 2 for the list of protein coding genes). Using these features, we first train an autoencoder model and then using the reduced feature set from the latent space of the autoencoder, we train a supervised feed-forward neural network that predicts healthy tissue versus tumor for each RNA-seq sample. The encoder in our autoencoder model has two latent layers followed by a third latent layer that produces the feature set with reduced dimensionality. The decoder mirrors the encoder. It takes in the features from the third latent layer of the encoder and reconstructs the gene expression values of the protein coding genes. Using the latent features from the encoder as input features, we then train a discriminator network with three latent layers followed by the output layer. Both the autoencoder and the discriminator networks use ReLU activation function and are trained using the adam optimizer (see Supplementary Table 3 for the detailed architecture of both models).

### LncRNA gene expression based model

For the next model, the input features are the gene expression of 14257 lncRNA genes. We extract the lncRNA genes from Ensembl BioMart, see Supplementary Table 4 for the list of lncRNA genes. The model architecture and training process of the lncRNA genes based model is similar to the protein-coding gene expression model described in the previous section (see Supplementary Table 5 for the detailed architecture of the lncRNA autoencoder and discriminator networks).

### Splicing junctions based model

Finally, the input features for our third model are the splicing quantification for 40147 alternative splice junctions from 11219 genes. We generated the splicing quantification for these junctions in the healthy tissues and tumors using MA-JIQ (see RNA-seq data processing section for details on quantification of these splicing junctions from the RNA-seq data). As we have thousands of splicing junctions as features, we again train an autoencoder for dimensionality reduction followed by a supervised neural network for tumor classification. The model architecture and training process of both these models is similar to the previous gene expression (protein-coding or lncRNA) based models (see Supplementary Table 6 for the detailed architecture of the splicing junctions based models).

### Interpretation of tumor classification models

In order to find the features responsible for classifying an RNA-seq sample as tumor, we employ the enhanced integrated gradients method for interpretation of a deep learning model. Here interpretation means attributing the prediction of a deep learning model to its input features. Enhanced integrated gradients computes feature attribution by aggregating gradients along a linear/non-linear path between a sample and a class-agnostic/specific baseline. Sample here refers to an input sample to the deep learning model. Baseline refers to model’s proxy to human counterfactual intuition. This implies that humans assign blame for difference in two entities on attributes that are present in one entity but absent in the other. Enhanced integrated gradients offers multiple baselines and paths. Since we want to find features that distinguish between tumor samples from healthy tissue samples, we use healthy tissues as the baseline class. Using median baseline from the healthy tissues class, we compute the attributions for the tumor samples by aggregating the gradients between the baseline and each tumor sample along a linear path in the original feature space. We assess class-wide significance of each feature by computing p-values by comparing the attribution distribution of a feature for the tumor class versus a random mixture of healthy tissues and tumors using a one-sided t-test with FDR correction for multiple hypothesis testing. For further details on enhanced integrated gradients please refer to (Jha et al., 2020).

### Selection of feature sets

High-attribution features were selected on the basis of their FDR-corrected (1%) p-value *<* 0.0001 and ranking above the knee-point of the curve in the case of positive attribution values or below the knee-point of the curve in the case of negative attribution values. A neutral set of size equivalent to the number of high-attribution features was selected among genes that had FDR-corrected p-value *>* 0.05 and ranking in the middle of the distribution of attribution values (attribution values close to 0). The list of COSMIC oncogenes and tumor suppressor genes (TSGs) was established from the COSMIC genes census that had a role in cancer comprising the annotation “oncogene” but not “TSG” for oncogenes and comprising the annotation “TSG” but not “oncogene” for TSGs.

### Gene ontology analysis, gene set enrichment analysis and Ingenuity Pathway analysis

Gene ontology analysis was performed with EnrichR (Chen et al., 2013) v.1.0 using a 2018 release of the GO Consortium annotations and including terms from the molecular function, cellular component and biological process categories. Gene set enrichment analysis was performed using GSEA4.1.0 (Mootha et al., 2003; Subramanian et al., 2005) against the msigdb.v7.4 gene set library and filtered for enrichment corresponding to hallmark, reactome and GO gene sets with a normalized p-value *<* 0.01.

### Functional characterizations

As for splice junction characterization, protein sequence used was translated from the sequence of the longest coding transcript in RefSeq. The conservation score was calculated from the summation of BLAST bit score from 6 species: human, chimpanzee, mouse, cattle, xenopus, zebrafish and chicken (taxids: 9606, 9598, 10090, 8364, 7955, 9031, 9913), normalized to the length of the human transcript. Loss of function mutation frequency is expressed as the gnomAD LOEUF score only for genes for which a LOEUF score was reported (Karczewski et al., 2020). Pyknon density was calculated using the list of human pyknons available from the pyknon database (Tsirigos & Rigoutsos, 2008) and shown as the number of pyknons found per 1,000 nt in the longest RefSeq annotated transcript.

## Acknowledgements

We would like to thank Joseph K. Aicher and Paul Jewell for their assistance in processing the gene expression and splicing data.

## Availability of data and materials

All processed data and code to reproduce figures will be available at https://bitbucket.org/biociphers/pan-cancer and be deposited in Zenodo before publication. RNA-Seq samples used for this study are listed in Supplementary Table 1.

## Author’s contributions

A.J. and M.Q.V. designed the study, analyzed and interpreted the data and wrote the manuscript; A.T.T, K.W.L. and Y.B. interpreted the data and wrote the manuscript.

---

**Figure S1.**
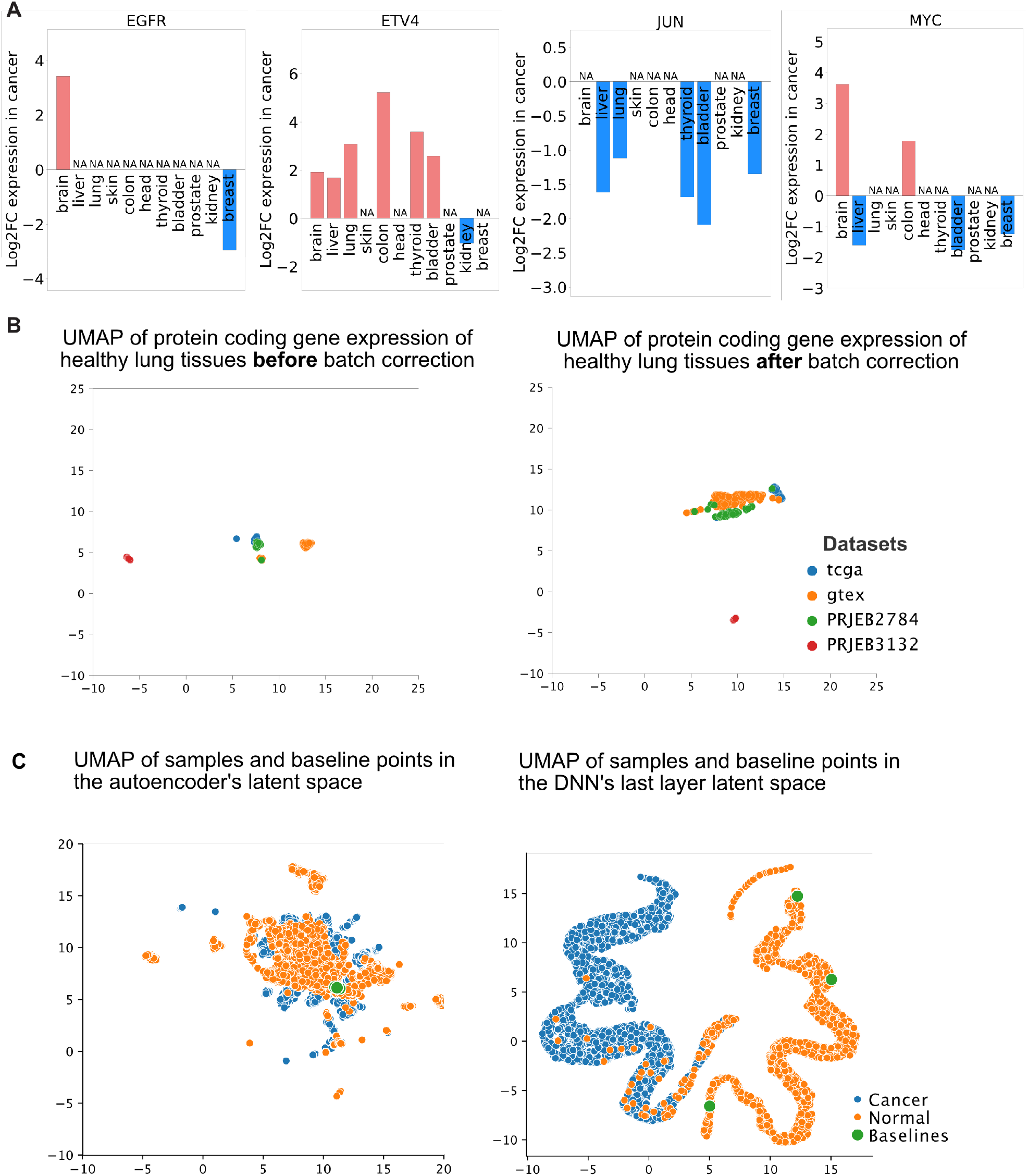
(A) Differential gene expression of four oncogenes in 11 healthy tissue-cancer pairs. NA: not differentially expressed. (B) UMAP showing the gene expression of protein coding genes in healthy lung samples in four datasets (GTEx, TCGA, PRJEB2784 and PRJEB3132) before (left) and after (right) batch correction. (C) Left: UMAP embedding of the protein-coding autoencoder’s latent layer representation of the cancer and normal samples along with the median baseline points from the normal class. Right: UMAP embedding of protein-coding DNN’s layer layer representation of the cancer and normal samples along with the median baseline points from the normal class.

**Figure S2.**
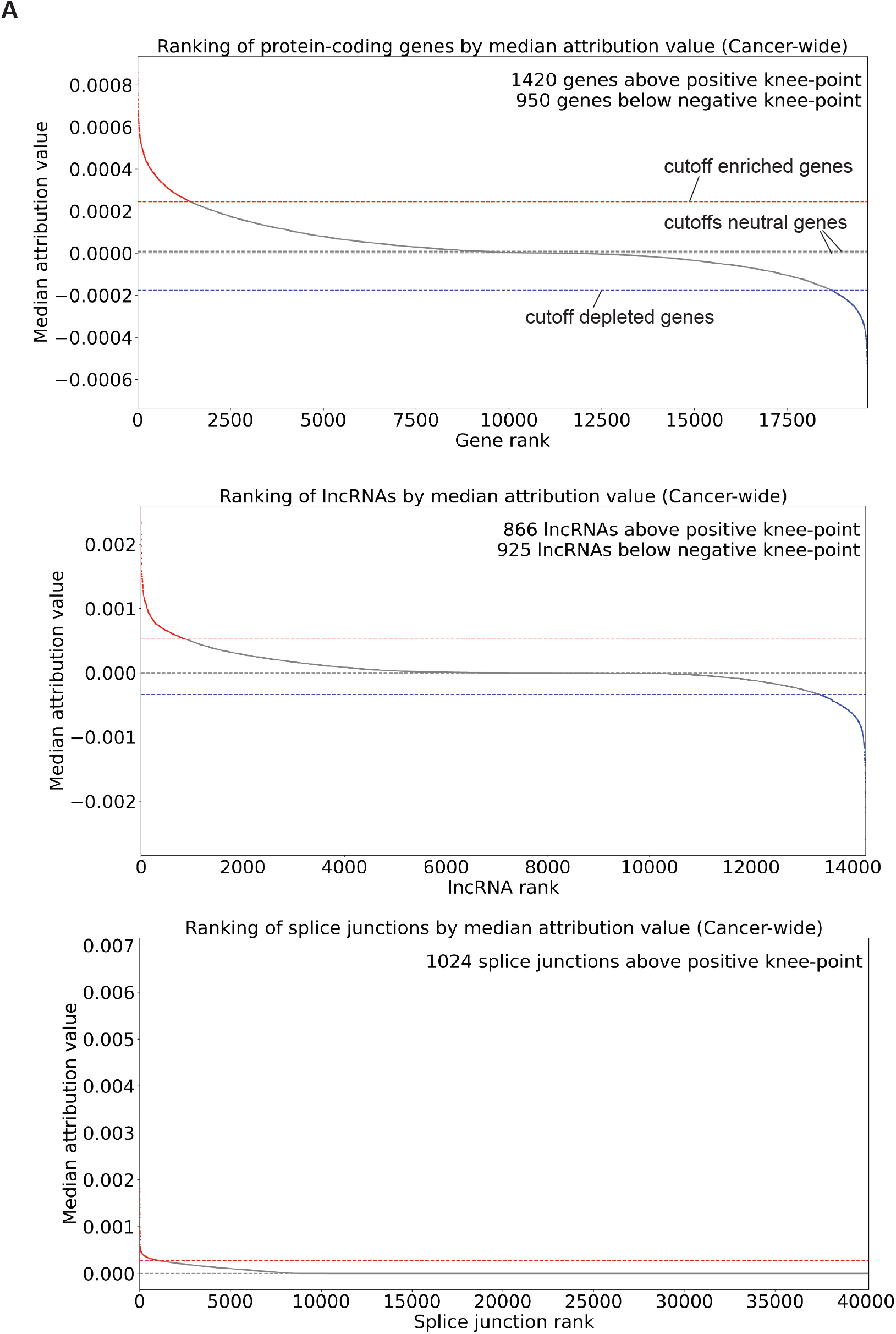
Distribution of transcriptomic features ranked from highest positive values to lowest negative values from models trained on protein-coding gene expression (top), lncRNA expression (middle) or splice junction usage data (bottom). Knee-point of the curves were determined using KneeLocator of the kneed package and are shown with broken horizontal lines.

**Figure S3.**
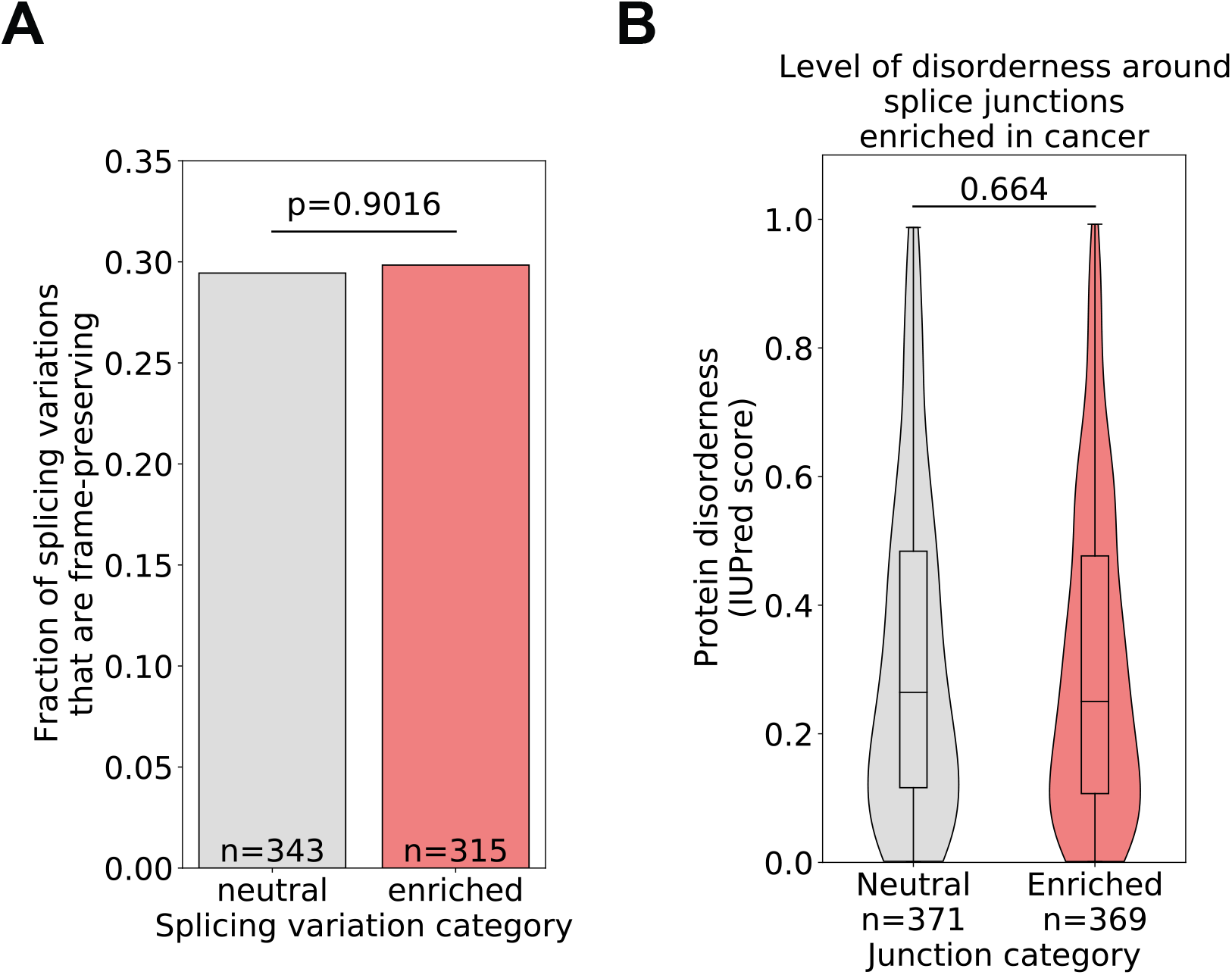
(A) Fraction of splicing variations where reading frame is preserved in both isoforms. (B) Predicted level of peptide disorderness around variable splice junctions.

**Figure S4.**
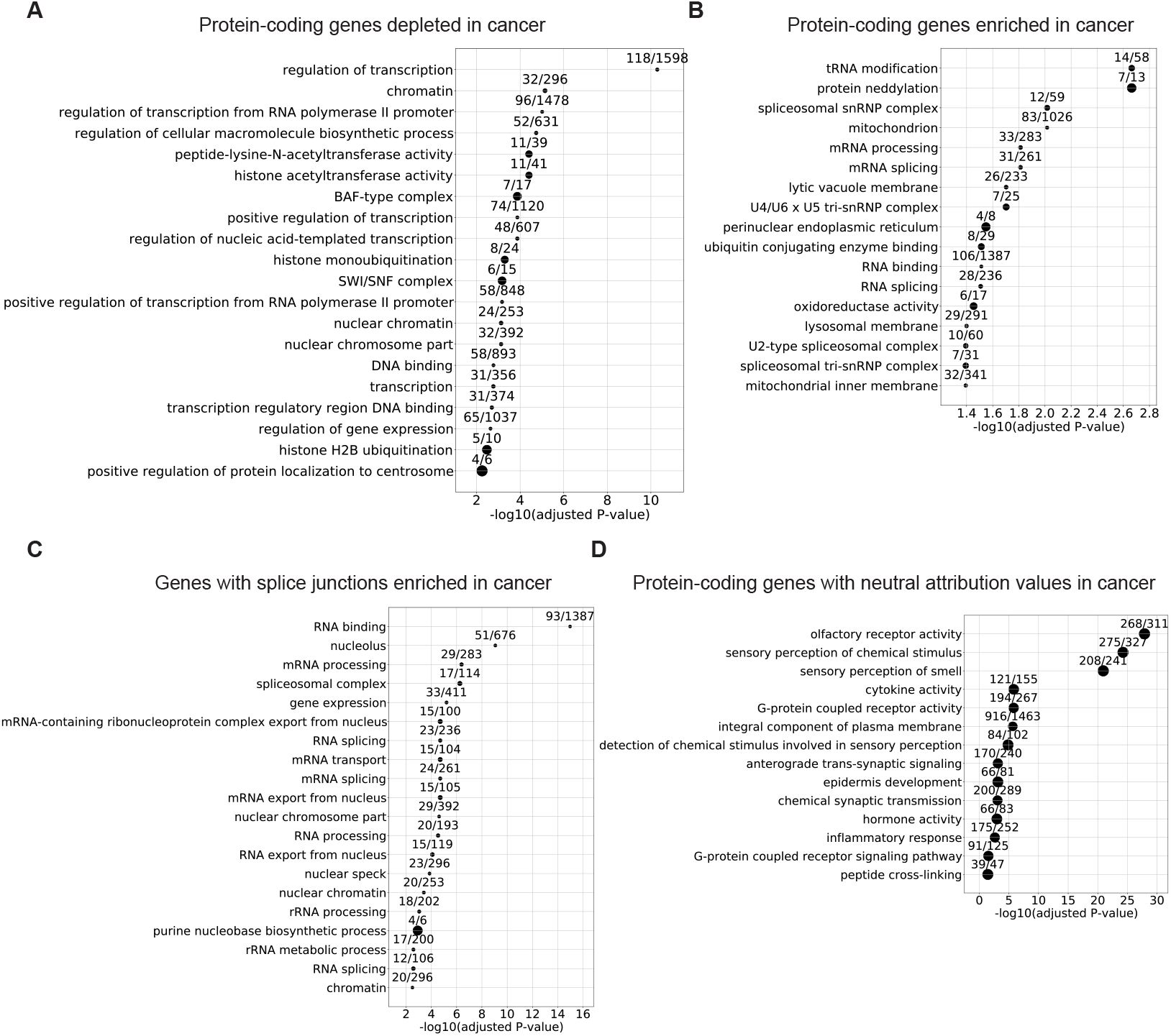
Gene ontology analysis of genes with high negative (A) or high positive (B) attribution values by expression, genes with splice junctions that have high attribution values (C), or genes with neutral attribution values by expression (D). Circle size and values indicate the fraction of genes corresponding to a GO term present in the gene set analyzed. Only top 20 terms are shown if more terms meet our cutoff of adjusted p-value *<* 0.05.

